# White matter microstructure in attention-deficit/hyperactivity disorder: a systematic tractography study in 654 individuals

**DOI:** 10.1101/787713

**Authors:** Christienne G. Damatac, Roselyne J. M. Chauvin, Marcel P. Zwiers, Daan van Rooij, Sophie E. A. Akkermans, Jilly Naaijen, Pieter J. Hoekstra, Catharina A. Hartman, Jaap Oosterlaan, Barbara Franke, Jan K. Buitelaar, Christian F. Beckmann, Emma Sprooten

**Affiliations:** Department of Cognitive Neuroscience, Donders Institute for Brain, Cognition and Behaviour, Radboud University Medical Center, Nijmegen, The Netherlands; Centre for Cognitive Neuroimaging, Donders Institute for Brain, Cognition and Behaviour, Radboud University, Nijmegen, The Netherlands; University of Groningen, University Medical Center Groningen, Department of Psychiatry, Groningen, The Netherlands; VU University Amsterdam, Department of Clinical Neuropsychology, Amsterdam, The Netherlands; Department of Human Genetics, Donders Institute for Brain, Cognition and Behaviour, Radboud University Medical Center, Nijmegen, The Netherlands; Department of Psychiatry, Donders Institute for Brain, Cognition and Behaviour, Radboud University Medical Center, Nijmegen, The Netherlands; Karakter Child and Adolescent Psychiatry University Centre, Nijmegen, The Netherlands; Centre for Functional MRI of the Brain, University of Oxford, Oxford, UK

**Author notes:** Correspondence Kapittelweg 29, 6525 EN Nijmegen, The Netherlands.

**Keywords:** ADHD, Diffusion MRI, Tractography, Neuroimaging, Dimensional, White matter

## Abstract

**Background:** Attention-deficit/hyperactivity disorder (ADHD) is a neurodevelopmental disorder characterized by age-inappropriate levels of inattention and/or hyperactivity-impulsivity (HI). ADHD has been related to differences in white matter (WM) microstructure. However, much remains unclear regarding the nature of these WM differences, and which clinical aspects of ADHD they reflect. We systematically investigated if FA is associated with current and/or lifetime categorical diagnosis, impairment in daily life, and continuous ADHD symptom measures.

**Methods:** Diffusion-weighted imaging (DWI) data were obtained from 654 participants (322 unaffected, 258 affected, 74 subthreshold; 7-29 years of age). We applied automated global probabilistic tractography on 18 major WM pathways. Linear mixed effects regression models were used to examine associations of clinical measures with overall brain and tract-specific fractional anisotropy (FA).

**Results:** There were significant interactions of tract with all ADHD variables on FA. There were no significant associations of FA with current or lifetime diagnosis, nor with impairment. Lower FA in the right cingulum’s angular bundle (rCAB) was associated with higher hyperactivity/impulsivity symptom severity (P_FWE_=0.045). There were no significant effects for other tracts.

**Conclusions:** This is the first time global probabilistic tractography has been applied to an ADHD dataset of this size. We found no evidence for altered FA in association with ADHD diagnosis. Our findings indicate that associations of FA with ADHD are not uniformly distributed across WM tracts. Continuous symptom measures of ADHD may be more sensitive to FA than diagnostic categories. The rCAB in particular may play a role in symptoms of hyperactivity and impulsivity.

## 1. Introduction

Attention-deficit/hyperactivity disorder (ADHD) is a common neurodevelopmental disorder characterized by age-inappropriate levels of inattention (IA) and/or hyperactivity-impulsivity (HI). Approximately 5% of children worldwide are diagnosed with ADHD and about 15% of youths with ADHD retain a full diagnosis, while around 70% retain impairing symptoms of the disorder in adulthood (1,2). Using diffusion-weighted imaging (DWI), studies have reported alterations in white matter (WM) microstructural properties across the lifespan of people with ADHD, some of which had an overlapping sample with the current study (3–11). However, generally limited sample sizes and methodological differences between studies contribute to inconsistencies in the locations and directions of findings thus far. Previous studies leave several unanswered questions that are critical for the assessment of the clinical relevance of such findings, especially whether WM microstructural properties are associated with: (1) trait versus state effects in ADHD; (2) continuous symptom measures versus categorical diagnosis of ADHD; and (3) clinical impairment in ADHD. Here, we applied automated tractography to DWI data from a large cohort of participants with and without ADHD to address such questions.

DWI measures the magnitude and direction of water diffusion, which in WM reflects the underlying organization of axons and their surrounding myelin (12–14). With diffusion tensor analysis, different metrics of water diffusion are calculated in each voxel, such as the degree of its directional preference (fractional anisotropy; FA), which is the most commonly investigated metric and the one we focus on here. Although FA is not a direct measure of physiological cellular properties, it is assumed to reflect a combination of the degree of parallel organization of axons, their packing density, and the amount and integrity of the surrounding myelin sheath (15,16). To date, findings from DWI in ADHD case-control studies have been largely mixed. Lower FA in ADHD has been found in a plethora of locations, most frequently in interhemispheric, frontal and temporal regions; yet, elevated FA has also been reported (3). Thus, many case-control studies show some differences in FA in ADHD, but the nature and anatomical locations of these findings are inconsistent.

FA has mostly been related to diagnosis at a single time-point, typically at the time of DWI acquisition. However, remission occurs in some patients, as do fluctuations in symptom severity and impairment over time (2). Typical case-control designs cannot disentangle stable lifetime trait effects associated with ADHD from those associated with ADHD as a current state. A better understanding of these dynamics has implications for assumptions behind genetic liability for ADHD and for the nature of the neural mechanisms underlying ADHD, and their potential receptiveness to treatment. Here, we refer to associations based on patients’ current diagnoses as “state” effects, and to associations related to ever-affected individuals regardless of current diagnosis as “trait” effects. Trait effects remain identifiable in remittent patients, pointing to the possibility that the diagnosis of ADHD at any point leaves an indelible ‘mark’ on WM that persists throughout life, regardless of the disorder’s progression (10). Longitudinal studies have found trait effects irrespective of diagnostic outcome: decreased FA has been found as a trait effect of ADHD in several thalamocortical tracts and the superior longitudinal fasciculus. However, in other studies, no such ADHD trait effects have been found (10,17). Larger sample sizes combined with sophisticated DWI analytical techniques may clarify inconsistencies in these previous findings.

Unlike the hard line of diagnostic category, ADHD symptoms are continually distributed throughout the population and the boundary between those with and without the disorder is ill-defined (18–20). Considering a continuous variable of symptom severity, rather than a categorical diagnosis, could increase power to detect ADHD-related cognitive processes and brain traits that are also continuously distributed. Non-clinical “control” participants may also exhibit subclinical ADHD characteristics and within patient groups there is considerable variation in symptom severity. Dimensional analyses allow for modeling of the entire spectrum of ADHD, from minor subthreshold symptoms to clinically extreme symptoms (21). Generally, however, DWI studies that have applied dimensional analyses have been inconsistent in terms of anatomical location and have suggested that, within the patient population, increased symptom severity ratings are associated with higher FA in widespread brain regions (9,22). Given that the categorical diagnosis of ADHD has generally been associated with reduced FA, a more in-depth study of these counterintuitive effects is warranted.

The sole presence of (sufficient) symptoms does not constitute a diagnosis of ADHD. For a clinical diagnosis, symptoms must be accompanied by impairment in daily functioning at home, at school or work, and/or in social settings. The degree of impairment does not directly map onto diagnosis or symptom severity scores. To understand the nature of case-control differences and their contrast with associations with symptom dimensions, an understanding of how clinical impairment is associated with brain differences is equally necessary. Although impairment may be a more subjective measure than symptom criteria, it is arguably the most impactful factor in the quality of life of patients. Impairment was found to be predictive of emotional lability in ADHD, independent of symptom severity (23). Magnetic resonance imaging (MRI) studies that considered clinical impairment as a separate factor, independent of diagnosis or symptom severity, are scarce to date.

An additional origin of discrepancies between studies may be found in methodological differences. The use of voxel-wise analyses in the presence of crossing fibers and anatomical differences in tract width and shape can lead to ambiguity in the anatomical location of DWI findings. In addition, residual effects of head motion after realignment tend to be associated with the main DWI outcome measures, but are not always taken into account (3). Previous tractography methods required user interaction (e.g. manually draw regions of interest or set thresholds for path angle and length), *a priori* selection of tracts, and employed local tractography (24). In an effort to improve large dataset analysis in a data-driven manner, we applied an automated tractography method to one of the largest cohorts of individuals with ADHD and healthy controls. We applied global probabilistic tractography in combination with anatomical knowledge of the tract’s location and shape, allowing fully automated tractography of several major WM tracts. In summary, in the largest DWI analysis of ADHD to date, we investigated the relationship of WM microstructure with lifetime and current diagnosis of ADHD, categorical and dimensional scales of current symptoms, and ADHD-related impairment.

## 2. Methods and Materials

### 2.1 Participants

A full description of the study design is available in previous work (25). Subjects were part of the International Multicenter ADHD Genetics (IMAGE; W1) study, which began in 2003 and included participants originally recruited with an ADHD diagnosis (probands), their affected and unaffected siblings, and healthy controls, as described previously (26). Data were collected at two centers: VU Amsterdam and Radboudumc Nijmegen. After a mean follow-up period of 5.9 years (SD=0.6), all W1 participants were invited for follow-up measurement. This second assessment wave (NeuroIMAGE1; W2) followed a phenotypic protocol similar to that of W1, but with the additional acquisition of MRI brain scans (25). A third assessment (NeuroIMAGE2; W3), occurred only in Nijmegen after approximately 3.7 years (SD=0.6) and followed the same protocol as that of W2.

The sample for our current analyses (summarized in Table 1) includes all individuals from W2 who had DWI scans that passed all quality control (n=570). We additionally included participants who had been newly recruited as part of W3 and thus had data available from only one wave (n=84 after all quality control). There were thus 654 participants from 366 families in total in the current analysis (age range: 7.72-28.59 years, mean age=17.41 years). Of these, 322 were unaffected, 258 had a full current diagnosis of ADHD, and 74 had a diagnosis of subthreshold ADHD at the first MRI acquisition time, as defined below. There were no differences on measures of ADHD severity (*P*=0.941), age (*P*=0.254), and sex (p=0.165) between subjects in the current analysis and the whole W2 sample (n=1085), including those who did not have available MRI data.

**Table 1.** Demographic and clinical characteristics of the ADHD affected, subthreshold, and unaffected groups based on participants’ diagnosis at the time of MRI acquisition.

|  | Affected<br>N = 258 |  | Subthreshold<br>N = 74 |  | Unaffected<br>N = 322 |  | Test Statistic | (P) |
| --- | --- | --- | --- | --- | --- | --- | --- | --- |
|  | Mean | (SD) | Mean | (SD) | Mean | (SD) |  |  |
| Age (years) | 17.4 | (3.6) | 18.2 | (3.6) | 17.2 | (3.7) | F(1,658) = 0.76 | (0.38) |
| IQ estimate | 95.1 | (16.0) | 102.2 | (13.5) | 104.3 | (14.9) | F(2,653) = 23.9 | (< 10<sup>-10</sup>) |
| Handedness (right) | 85% | N = 219 | 82% | N = 61 | 86% | N = 276 | $\chi^2(4) = 1.1$ | (0.89) |
| Sex (female) | 30% | N = 76 | 42% | N = 31 | 53% | N = 170 | $\chi^2(2) = 34.6$ | (< 10<sup>-8</sup>) |
| Scan site (Nijmegen) | 57% | N = 148 | 59% | N = 44 | 52% | N = 167 | $\chi^2(2) = 5.1$ | (0.08) |
| Head motion | 0.76 | (1.1) | 0.75 | (1.3) | 0.59 | (0.4) | F(2,655) = 3.1 | (0.05) |
| Diagnosed ever (yes) | 100% | N = 258 | 32% | N = 24 | 6.2% | N = 20 |  |  |
| CPRS HI + IA | 22.4 | (12.1) | 11.1 | (9.6) | 4.3 | (4.9) |  |  |
| CPRS HI | 9.6 | (6.5) | 3.7 | (4.5) | 1.3 | (2.1) |  |  |
| CPRS IA | 14.2 | (7.1) | 7.2 | (5.6) | 2.9 | (3.5) |  |  |
| K-SADS-PL HI | 5.2 | (2.5) | 2.8 | (2.0) | 2.1 | (2.0) |  |  |
| K-SADS-PL IA | 7.0 | (1.4) | 4.7 | (0.8) | 6.0 | (2.1) |  |  |
| Impaired (yes) | 98% | N = 252 | 47% | N = 35 | 3.7% | N = 12 |  |  |
| Duration medication use (days) | 1413 | (332) | 645 | (1111) | 46 | (332) |  |  |
| History of medication use (yes) | 78% | N = 202 | 36% | N = 27 | 3.1% | N = 10 |  |  |
| *Comorbidity* |  |  |  |  |  |  |  |  |
| Conduct disorder | 4.7% | N = 12 | 1.4% | N = 1 | 0% | N = 0 | $\chi^2(2) = 1.3$ | (0.53) |
| Major depression | 0.8% | N = 2 | 0% | N = 0 | 0.3% | N = 1 | $\chi^2(2) = 1.3$ | (0.52) |
| Oppositional defiant disorder | 30% | N = 76 | 8.1% | N = 6 | 1.6% | N = 5 | $\chi^2(2) = 93.5$ | (< 10<sup>-15</sup>) |
| *Substance use ever* |  |  |  |  |  |  |  |  |
| Alcohol | 20% | N = 52 | 18% | N = 13 | 16% | N = 51 | $\chi^2(2) = 1.7$ | (0.42) |
| Tobacco | 48% | N = 125 | 36% | N = 27 | 32% | N = 102 | $\chi^2(2) = 16.8$ | (0.0002) |
| Cannabis or hash | 28% | N = 72 | 24% | N = 18 | 18% | N = 57 | $\chi^2(2) = 8.9$ | (0.01) |
| Other | 10% | N = 26 | 9.5% | N = 7 | 4.7% | N = 15 | $\chi^2(2) = 6.9$ | (0.03) |

Demographic between-group differences were tested using F-tests for continuous variables and $\chi^2$ -tests for categorical variables. Totals are for all participants who were included in the final sample, after all quality control.
Medications: Ritalin (methylphenidate), Concerta (methylphenidate), Strattera (atomoxetine), and any other ADHD medication.
CPRS: Conners Parent Rating Scale
HI: hyperactivity-impulsivity
IA: inattention

### 2.2 Clinical assessments

A full description of the clinical assessments in our sample is available in previous work (25). ADHD categorical diagnosis, clinical impairment, and symptom dimension severity scores for IA and HI were determined through the Kiddie – Schedule for Affective Disorder and Schizophrenia Present and Lifetime Version (K-SADS-PL) and the Conners Parent Rating Scale (CPRS) questionnaires at W1, W2, and W3 (27,28). An algorithm was applied to create a combined symptom count from the interview and questionnaires, as detailed previously (25). A participant <18 years was diagnosed with ADHD according to DSM-IV criteria (if he/she had ≥6 IA and/or ≥6 HI symptoms; or ≥5 symptoms if participant was ≥18 years old), causing functional impairment at home, school or work, or in social settings (29). All unaffected participants (including unaffected siblings and unrelated controls) were required to have a score of ≤3 in both symptom dimensions (≤2 symptoms, if participant was ≥18 years old). Participants who fulfilled criteria for neither ADHD nor unaffected status were classified as subthreshold ADHD. Besides diagnosis and symptom score, multiple setting clinical impairment was investigated separately, defined as impairment in at least 2 of the 3 domains according to the K-SADS-PL interview: school/work, home, and social.

The majority of patients were taking prescription medication for ADHD, mostly methylphenidate or atomoxetine. The duration of medication use was recorded as the cumulative number of days of use, while the history of medication use was recorded as whether or not the participant had ever taken ADHD medication. Further details on medication use in our sample is available in previous work (30,31). A history of comorbid disorders was screened for via the K-SADS-PL semi-structured interview (27,32). For children <12 years old, the child’s parents or researchers assisted in the completion of the self-report questionnaires. Participants with elevated scores on one or more of the K-SADS screening questions were further asked to complete a full supplement for each disorder. The final diagnosis was based on DSM-IV criteria of each disorder. Lastly, alcohol use, nicotine use, and other drug use at any point before assessment was recorded through self-report (33–35).

### 2.3 Imaging acquisition and analysis

MRI data were acquired with either a 1.5-Tesla MAGNETOM-Sonata or a 1.5-Tesla AVANTO scanner (Siemens, Erlangen, Germany). Both scanners were equipped with the same 8-channel phased-array head coil. Details of the T1-weighted and DWI data acquisition have been described previously (25). DWI images were preprocessed and included motion and eddy-current corrections (25,36). Diffusion tensor eigenvectors, eigenvalues, and FA were then calculated for each voxel (37). Bedpostx was applied with a two-fiber model to estimate the distributions of the diffusion parameters for probabilistic tractography (38).

We then applied Tracts Constrained by Underlying Anatomy (TRACULA; 24; http://surfer.nmr.mgh.harvard.edu/fswiki/Tracula), a FreeSurfer toolbox, to delineate 18 major WM tracts (Figure 1). TRACULA is a method for the automated reconstruction of major WM pathways based on an earlier well-known global probabilistic approach (39). TRACULA extends this algorithm by incorporating anatomical knowledge in the prior probability function, so that the final segmented tract is not only the best fit given the observed diffusion data within each subject, but also given its similarity to the known tract anatomy in relation to grey matter segmentations from FreeSurfer.

**Figure 1.**
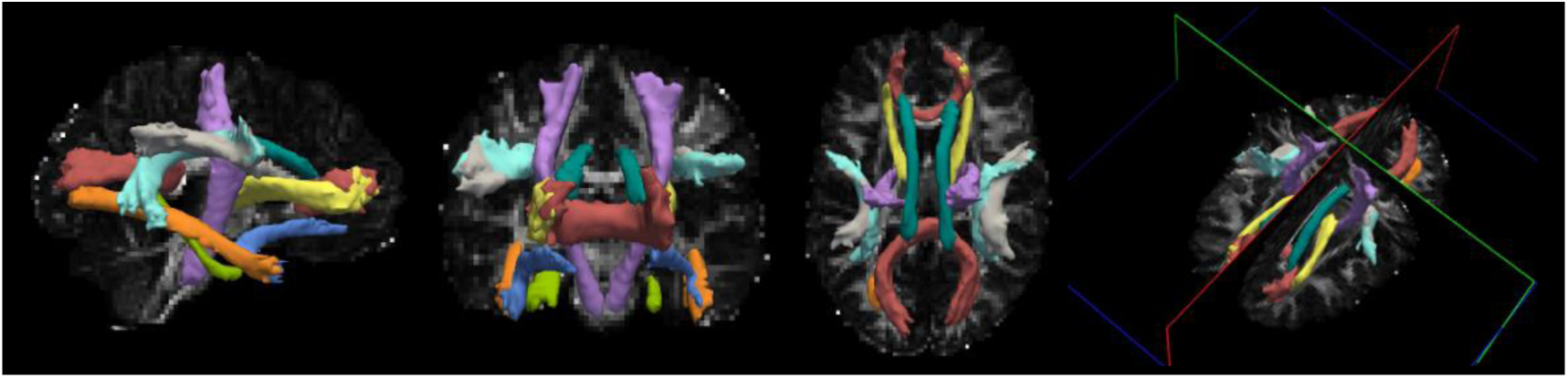
Example pathway reconstruction and merged 4D volume output of TRACULA in a healthy control subject. All 18 white matter tracts (8 bilateral and 2 interhemispheric) are displayed at 20% of their maximum probability distribution threshold as an isosurface over the subject’s FreeSurfer segmentation: 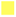 anterior thalamic radiations (ATR), 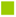 cingulum-angular bundle (CAB), 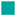 cingulum-cingulate gyrus bundle (CCG), 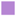 corticospinal tract (CST), 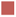 corpus callosum forceps major and minor (FMAJ, FMIN), 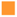 inferior longitudinal fasciculus (ILF), 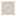 superior longitudinal fasciculus-parietal terminations (SLFP), 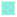 superior longitudinal fasciculus-temporal terminations (SLFT), 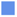 uncinate fasciculus (UNC).

We used FreeSurfer 5.3 to define cortical and subcortical regions in the T1-weighted images of each individual (40). FreeSurfer segmentations were checked for quality based on a modified version of the ENIGMA Protocol 2.0 (http://enigma.ini.usc.edu/protocols/imaging-protocols/), resulting in n=22 exclusions. Next, ‘trac-all’ was run in TRACULA to segment all of the above-mentioned tracts in native space. We visually checked the anatomical accuracy of each subject’s tract segmentation output, resulting in n=7 exclusions (https://surfer.nmr.mgh.harvard.edu/fswiki/FreeviewGuide). See Figure 1 for an example of TRACULA’s segmentation output in a single healthy control subject. Finally, each participant’s scan was registered to MNI space for analyses of FA at each location along the tract.

### 2.4 Statistical analyses

All statistical analyses were performed in R3.4.2(https://cran.r-project.org/). We investigated 4 types of effects and tested each separately as a fixed factor: (1) current ADHD diagnosis at the time of first MRI acquisition (state), (2) having ever been diagnosed with ADHD (trait), (3) clinical impairment, and (4) total symptom scores on HI and IA dimensions separately. In categorical analyses 1 and 2, the subthreshold group was excluded. In analysis 1: 322 participants were unaffected and 258 were affected with ADHD. For analysis 2: 312 participants had not been diagnosed with ADHD by the time of scan (never), while 292 had been diagnosed by the time of scan (ever).

For all analyses, a step-wise approach was taken, where first global effects across all tracts and interactions with tract were tested. *Only* if we found a significant overall effect or a significant interaction effect with tract, we examined the effect of the independent variable in question on FA in individual tracts. For these tract-specific analyses, *P*-values were Bonferroni-corrected for the number of tracts, reported as *P*_FWE_ (*P*_FWE_=*P*x18). For both global and by-tract analyses, we applied linear mixed effects regression models (R package ‘lme4’ version 1.1-21). All analyses incorporated age, sex, MRI acquisition site, assessment wave, and head motion (framewise displacement) as fixed effects and family membership as a random effect. Global models also included tract as a fixed effect and subject as a random effect. See <u>Table S7</u> for a summary of our models.

If any tract was significant in the previous analyses, we performed a point-wise comparison of FA per voxel for every location along the tract.

In our post hoc analyses, we investigated effects of medication use by separately adding use of any ADHD medication as a binary factor and then as a continuous factor. Potential effects of IQ, substance use and major comorbidities were also separately analyzed through the addition of each variable as a fixed factor.

## 3. Results

### 3.1 State and trait effects of ADHD diagnosis

There was no association of ADHD state or ADHD trait with FA, globally across all tracts (Table 2). Both ADHD state and trait showed highly significant interaction effects with tract (*P*<0.0006), which led us to further examine FA differences by tract (tract-specific models). However, there were no significant effects on FA for any specific tract; in fact, all uncorrected P-values were >0.05 (Tables S2, S3).

**Table 2.** All global model analyses.

| | *t* | *df* | Std $\beta$ | *P* | Tract interaction *P* |
| --- | --- | --- | --- | --- | --- |
| State | -0.924 | 489 | -0.011 | 0.356 | **0.000537** |
| Trait | -0.513 | 584 | -0.006 | 0.608 | **0.000003** |
| Impairment | -1.337 | 571 | -0.016 | 0.182 | **0.000020** |
| CPRS HI + IA | -2.280 | 567 | -0.027 | **0.023** | **0.000140** |
| CPRS HI | -2.160 | 556 | -0.026 | **0.031** | **0.000017** |
| CPRS IA | -2.174 | 564 | -0.026 | **0.030** | **0.006162** |

### 3.2 Clinical impairment

Our global model for the association between FA and impairment resulted in no association with impairment and an interaction effect with tract (*P*=0.00002). No tracts had a significant association of FA with impairment (Table S3).

### 3.3 Continuous symptom scores

The global models indicated main effects of IA and HI CPRS scores (respectively *P*=0.030 and *P*=0.031) and again significant interaction effects with tract (Both *P*<0.007) (Table 2). Tract-specific models for HI CPRS score showed a negative association between rCAB FA and HI (*P*_*FWE*_=0.045,*t*=-3.04,*df*=632). Figure 3 provides a closer inspection of this effect in the rCAB for each symptom dimension against FA. Other tracts did not show associations between FA and either symptom dimension at Bonferroni-corrected significance levels (Figure 2 and Table S4).

**Figure 2.**
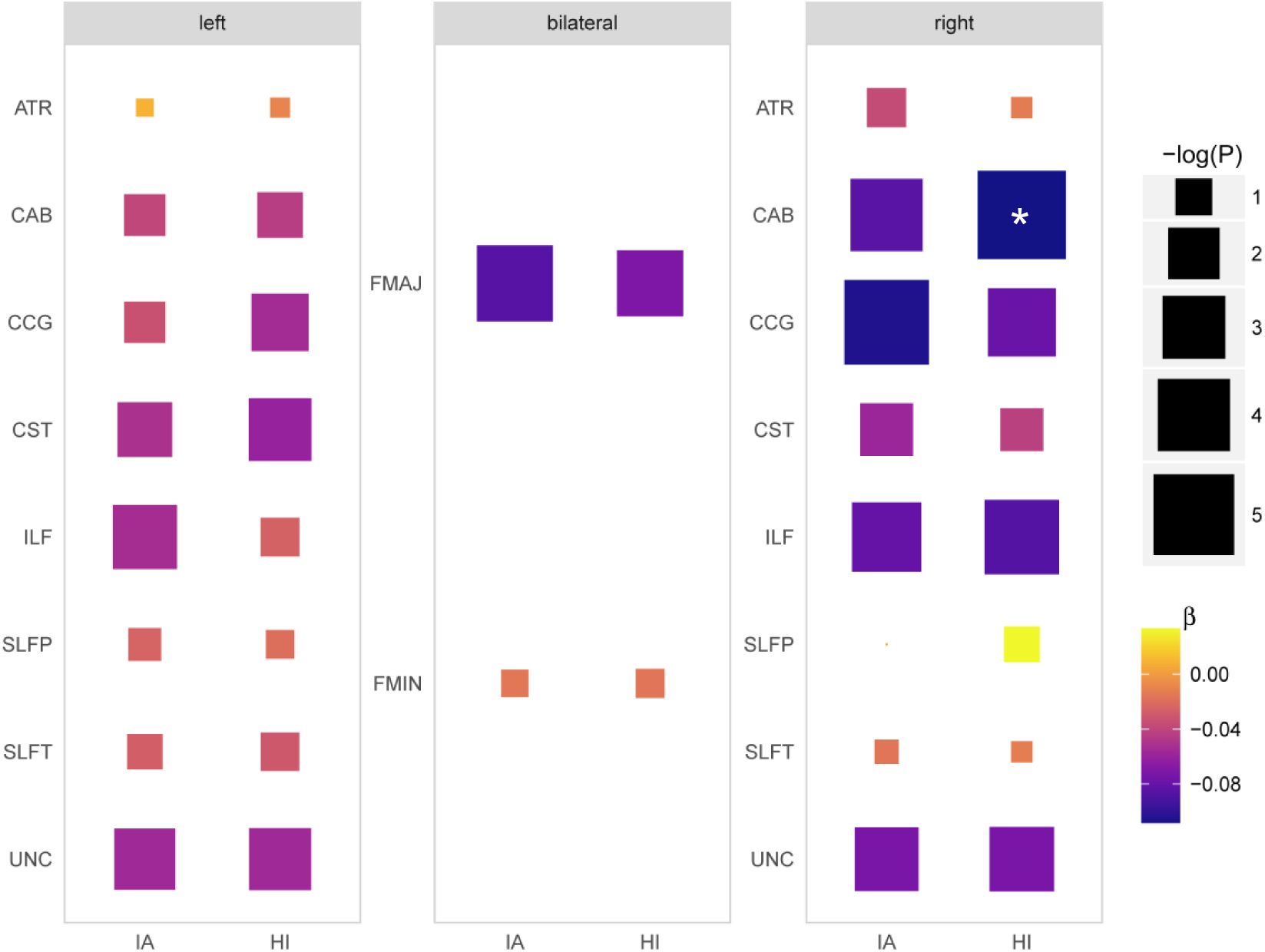
All tract-specific models for the effect of CPRS symptom dimension (IA and HI) on FA. Uncorrected *P*-values and standardized ß coefficients: negative log of the *P*-values are represented by size so that more significant values are larger; more negative ß coefficients are displayed in darker color; the only significant tract-specific model is marked by an asterisk. See Table S4 for associated statistics.

**Figure 3.**
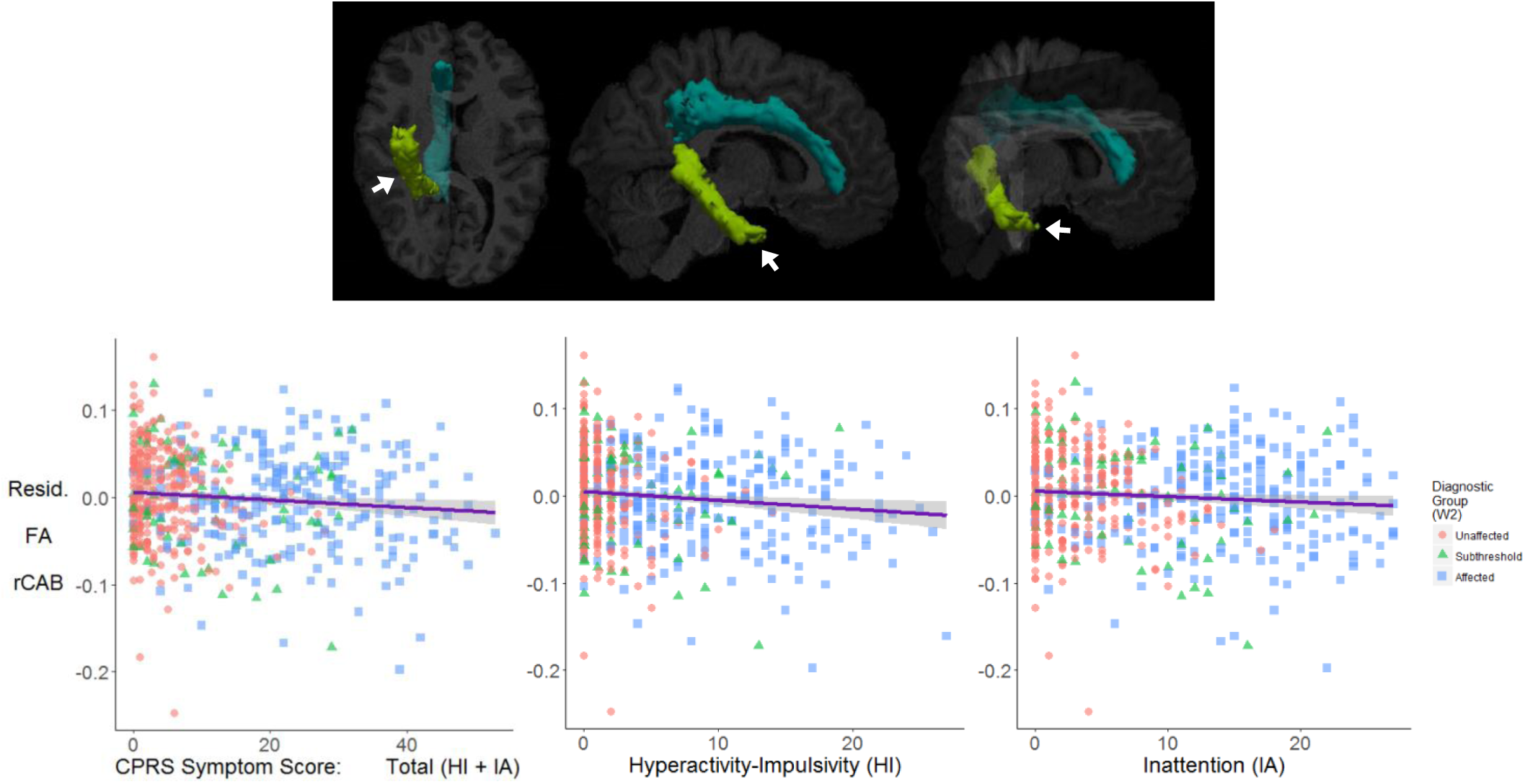
Tract-specific rCAB mean FA and symptom score. Top: TRACULA reconstruction in a healthy control of the rCAB in green and indicated by white arrows, from inferior, right and ventral anterior views, respectively. The cingulum’s cingulate gyrus bundle is shown in blue for reference. Bottom: Scatterplots with regression lines (95% confidence intervals) of the association between each dimension score (combined, HI, and IA) and the residuals of rCAB mean FA. Data points are grouped according to each participant’s diagnosis at W2: 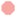 Unaffected, 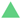 Subthreshold, 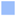 Affected. Mean FA in the rCAB had a negative association with HI symptom score, but not IA. See Table S4 for associated statistics.

### 3.4 Point-wise comparison of FA along rCAB

The effect of IA and HI scores on FA was relatively equally distributed along the length of the tract, with slightly larger negative effects at the posterior end compared to the anterior end (Figure 4). There is no particular rCAB area or voxel that contributed significantly more than others to the association between FA and HI score, nor to that of FA and IA score (Table S5).

**Figure 4.**
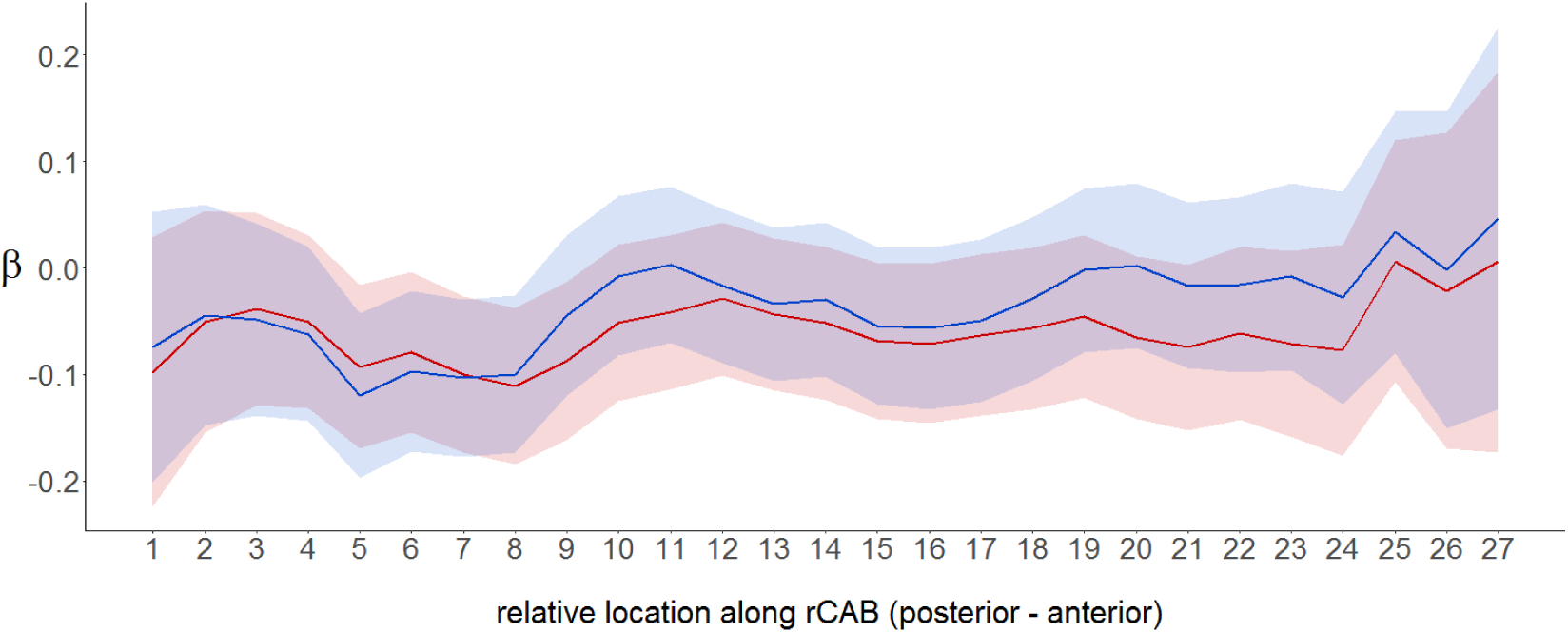
Point-wise comparison of FA along rCAB. Colored lines show the standardized ß coefficients (95% confidence intervals) of the associations between FA along the length of the rCAB and symptom score, grouped by symptom dimension, shown from posterior to anterior ends of the tract. ß coefficients of FA associations with HI symptom score are in red and those with IA are in blue for each of the 27 cross-sections along the standard-space tract spline. Towards each endpoint of the tract, fewer participants contributed FA data to the analysis, reflected in wider confidence intervals at the tract ends. See Table S5 for associated statistics.

### 3.5 Post hoc analyses

Correlations between global and rCAB FA and medication use, comorbid disorders, substance use, and IQ are presented in Table 3. Days of medication use was nominally associated with rCAB FA (*P*=0.042). Therefore, to test for potential confounding effects of medication use duration, we added it as a covariate to the previously significant model of HI on FA in the rCAB. When medication use duration was included as a covariate, the effect of HI was no longer significant at our Bonferroni-corrected significance level (*P*=0.028,*t*=-2.207,*df*=570), but neither was the effect of medication itself (*P*=0.222,*t*=-1.223,*df*=570). Medication use and HI symptom scores were highly collinear, given a correlation when cases and controls were pooled (Spearman ρ=0.661). Medication use did not have an interaction effect with HI scores (*P*>0.069).

**Table 3.** Post hoc analyses.

|  |  | Global FA |  |  | Tract-specific rCAB FA |  |  |
| --- | --- | --- | --- | --- | --- | --- | --- |
| | | Std $\beta$ | $t$ | $P$ | Std $\beta$ | $t$ | $P$ |
| Medication use | Binary (yes or no) | -0.003 | -0.206 | 0.837 | -0.033 | -0.845 | 0.399 |
|  | Continuous (days) | -0.010 | -0.803 | 0.422 | -0.078 | -2.039 | **0.042** |
| Comorbidities<br>(yes or no) | Conduct disorder | -0.037 | -0.714 | 0.480 | -0.004 | -0.116 | 0.908 |
|  | Major depression | 0.095 | 1.975 | 0.051 | -0.018 | -0.512 | 0.609 |
|  | Oppositional defiant disorder | -0.016 | -1.352 | 0.177 | -0.059 | -1.626 | 0.105 |
| Substance use<br>(yes or no) | Alcohol | -0.003 | -0.211 | 0.833 | -0.005 | -0.142 | 0.887 |
|  | Tobacco | 0.009 | 0.660 | 0.510 | 0.010 | 0.231 | 0.818 |
|  | Cannabis or hash | 0.002 | 0.118 | 0.906 | 0.052 | 1.252 | 0.211 |
|  | Other drugs | -0.010 | -0.719 | 0.473 | -0.012 | -0.313 | 0.754 |
| IQ estimate |  | -0.010 | -0.837 | 0.403 | 0.012 | 0.315 | 0.753 |

Standardized $\beta$ coefficients, associated $t$ and uncorrected $P$ values for the regression of medication use, comorbidities, and drug use until the date of MRI scan against global and rCAB FA. We found a negative effect of total duration of medication use as a continuous variable on FA, but not history of medication use as a binary factor. No other factors had a significant effect on either global or rCAB FA.

Head motion, age, and sex were already included as covariates in all models. There were no significant main or interaction effects of any of these variables with any independent variable of interest on FA for all analyses reported above. Although affected and subthreshold individuals displayed greater head motion than those who were unaffected, this difference was not significant (Table 1) and there was never an interaction effect with motion in all analyses.

## 4. Discussion

We performed a large, systematic analysis of ADHD clinical measures with WM microstructure using automated tractography. The presence of significant interactions with tract indicates that, for all ADHD-related variables tested, ADHD’s effect on FA varies across the different WM tracts. However, we did not find any evidence for FA reductions or increases in association with ADHD diagnosis, neither as trait nor state measure, nor with impairment, across 18 major white matter tracts. In contrast, regarding the HI symptom domain as a dimensional measure, our results showed reduced FA in the temporal portion of the cingulum bundle, or rCAB. Hence, our findings here suggest that continuous measures of current symptom severity state may be more sensitive to differences in FA in association with ADHD than are categorical clinical measures.

The cingulum has been previously implicated in voxel-based studies of ADHD, but with mixed exact locations and directions of effects (2,41,42). This may be due to differences in DWI analyses, wherein the cingulum is not easily delineated by various analytic methods. Only some connections encompass the entire length of the tract and discrete subdivisions of the cingulum display distinct FA measures at different medial-lateral positions within the bundle, indicating qualitative changes along the length of the tract (41–43). The present separation of the CAB from its dorsal counterpart may have provided greater resolution for identifying associations that were heretofore difficult to detect, or were manifested differently. Moreover, our rCAB point-wise analysis revealed that IA and HI symptom effects on FA are comparable for all voxels, which indicates that our result was driven by a relatively subtle effect across the entire tract—an effect that could be more challenging to detect with voxel-based or less sophisticated tractography methods.

The CAB links the posterior default mode network (DMN) and the medial temporal lobe. Aberrant functional connectivity involving the DMN is one of the most consistent neuroimaging characteristics of ADHD (44–48). Greater severity of HI and IA have also been associated with decreased DMN connectivity (49,50). The DMN is characterized by increased activation during rest and mind wandering and has also been associated with emotional lability (51–57). Mind wandering has been associated with emotional lability and greater ADHD symptom severity (58). Our finding of reduced FA in the rCAB could speculatively be an anatomical substrate of the frequently observed altered DMN functional connectivity in ADHD, and the associated emotional problems and increased mind wandering that may stem from aberrant activation in the DMN and its connections with the medial temporal lobe.

Given the current largest DWI ADHD sample yet and our use of a validated robust global probabilistic tractography method, the persistence of null results in association with ADHD diagnosis is noteworthy. We can speculate as to why our analyses did not generally replicate the rather large literature reporting WM alterations related to ADHD. Head motion can produce spurious DWI findings, which is of particular concern regarding ADHD given HI is associated with greater motion (59,60). A previous TRACULA analysis of group differences in DWI measures demonstrated that several tracts, including the rCAB, had greater group differences in DWI measures when group differences in head motion were higher (60). Until recently, many DWI studies of ADHD did not examine group differences in head motion and, similar to our categorical results here, most studies that reported no case-control difference in motion also had null results (3). All of our analyses included head motion as a covariate and each model was checked for interaction effects with head motion.

Days of medication use was negatively associated with rCAB FA and, when added into our rCAB model, resulted in a loss of the main effect of HI score on FA. This limitation is not surprising, given that— especially when cases and controls were pooled—ADHD medication use and symptom score were highly correlated. High levels of co-linearity between medication and symptom severity means that, within the context of this study design, we cannot clearly separate these effects. In contrast to our observations here, other studies have shown that methylphenidate treatment is associated with higher global FA, while other more suitably designed DTI studies that specifically investigated medication effects in ADHD patients did not find confounding effects of medication use on FA (61–63). The attenuated significance of both medication and HI symptoms when included in the same model tentatively supports an effect of ADHD symptomatology in combination with a secondary effect of medication use—merely due to higher medication use by those with more symptoms.

Finally, we must note that the physiological interpretation of DWI findings remains somewhat speculative (15,64). For example, in a single fiber bundle, decreased FA may represent myelin breakdown or reduced axonal integrity, whereas in regions with crossing fibers, it may represent increased neuronal branching and could, as such, even indicate increased structural connectivity. Hence, we must be cautious in interpreting our findings in terms of specific neurobiological mechanisms. In contrast to many previous studies that used TBSS or other voxel-based methods, we used tractography. This allows locating any FA differences to fiber tracts with known anatomy, which aids in relating it to brain networks and brain function. In addition, tractography allows for more inter-subject variation in the shape and size of fibers and the total brain, without affecting the FA values within the voxels. On the other hand, tractography is less sensitive to the very subtle but spatially consistent effects that are often reported using TBSS in combination with threshold-free cluster enhancement (65). Thus, differences in spatial location and extent between our study and previous DWI studies in ADHD should be interpreted bearing in mind these methodological choices.

### 4.1 Conclusion

In conclusion, to our knowledge, this is the first time global probabilistic tractography has been applied to an ADHD DWI dataset of this size. Using TRACULA allowed for the automatic, simultaneous extraction of FA measures from hundreds of subjects. In line with previous data, we provide further evidence that accentuates the complexity of this disorder and complements information from case-control studies. We found that FA is associated with symptom dimension scores, but not with categorical diagnostic ADHD measures. We conclude that associations of FA with ADHD may be more subtle than previously thought, and that the rCAB may be more involved in the symptoms of ADHD than previously thought.

## 7. Supplement

**Table S1.** State tract-specific model analyses.

| Hemisphere | Tract | *t* | df | Std $\beta$ | *P* | *P<sub>FWE</sub>* |
| --- | --- | --- | --- | --- | --- | --- |
| Left | ATR | 0.65 | 552 | 0.021 | 0.516 | 1 |
|  | CAB | -0.72 | 571 | -0.027 | 0.472 | 1 |
|  | CCG | 1.10 | 571 | 0.037 | 0.270 | 1 |
|  | CST | -1.22 | 556 | -0.041 | 0.223 | 1 |
|  | ILF | -0.64 | 548 | -0.018 | 0.520 | 1 |
|  | SLFP | 0.03 | 552 | 0.001 | 0.974 | 1 |
|  | SLFT | -0.21 | 549 | -0.007 | 0.833 | 1 |
|  | UNC | -0.36 | 553 | -0.011 | 0.716 | 1 |
| Bilateral | FMAJ | -1.67 | 564 | -0.061 | 0.096 | 1 |
|  | FMIN | 0.51 | 537 | 0.014 | 0.607 | 1 |
| Right | ATR | 0.28 | 568 | 0.011 | 0.776 | 1 |
|  | CAB | -1.61 | 574 | -0.062 | 0.108 | 1 |
|  | CCG | -0.80 | 569 | -0.031 | 0.422 | 1 |
|  | CST | -0.69 | 567 | -0.028 | 0.494 | 1 |
|  | ILF | -1.90 | 562 | -0.072 | 0.058 | 1 |
|  | SLFP | 1.04 | 559 | 0.042 | 0.298 | 1 |
|  | SLFT | 0.25 | 548 | 0.010 | 0.799 | 1 |
|  | UNC | -0.17 | 550 | -0.007 | 0.863 | 1 |

Associated *t*-values, degrees of freedom, standardized beta coefficients, uncorrected *P*-values, and interaction with tract *P*-values wherein ADHD was modeled as a state.

**Table S2.** Trait tract-specific model analyses.

| Hemisphere | Tract | *t* | df | Std $\beta$ | *P* | *P<sub>FWE</sub>* |
| --- | --- | --- | --- | --- | --- | --- |
| Left | ATR | 1.43 | 620 | 0.044 | 0.154 | 1 |
|  | CAB | -1.09 | 647 | -0.039 | 0.277 | 1 |
|  | CCG | 1.54 | 646 | 0.048 | 0.125 | 1 |
|  | CST | -0.71 | 627 | -0.022 | 0.481 | 1 |
|  | ILF | -0.72 | 612 | -0.019 | 0.472 | 1 |
|  | SLFP | 0.13 | 633 | 0.004 | 0.900 | 1 |
|  | SLFT | -0.23 | 627 | -0.007 | 0.815 | 1 |
|  | UNC | -0.53 | 619 | -0.016 | 0.593 | 1 |
| Bilateral | FMAJ | -0.92 | 640 | -0.031 | 0.360 | 1 |
|  | FMIN | 1.04 | 626 | 0.029 | 0.300 | 1 |
| Right | ATR | 0.28 | 638 | 0.010 | 0.779 | 1 |
|  | CAB | -1.66 | 649 | -0.060 | 0.097 | 1 |
|  | CCG | -0.50 | 640 | -0.018 | 0.616 | 1 |
|  | CST | 0.34 | 638 | 0.013 | 0.737 | 1 |
|  | ILF | -1.71 | 638 | -0.062 | 0.088 | 1 |
|  | SLFP | 1.18 | 634 | 0.044 | 0.239 | 1 |
|  | SLFT | 0.43 | 616 | 0.015 | 0.664 | 1 |
|  | UNC | -0.18 | 625 | -0.007 | 0.856 | 1 |

Associated *t*-values, degrees of freedom, standardized beta coefficients, uncorrected *P*-values, and interaction with tract *P*-values for trait.

**Table S3.** Impairment tract-specific model analyses.

| Hemisphere | Tract | *t* | df | Std $\beta$ | *P* | *P<sub>FWE</sub>* |
| --- | --- | --- | --- | --- | --- | --- |
| Left | ATR | 1.12 | 608 | 0.034 | 0.264 | 1 |
|  | CAB | -1.09 | 639 | -0.039 | 0.278 | 1 |
|  | CCG | 1.45 | 637 | 0.046 | 0.146 | 1 |
|  | CST | -1.45 | 617 | -0.046 | 0.147 | 1 |
|  | ILF | -1.31 | 602 | -0.035 | 0.190 | 1 |
|  | SLFP | -0.16 | 624 | -0.005 | 0.874 | 1 |
|  | SLFT | -0.43 | 618 | -0.014 | 0.670 | 1 |
|  | UNC | -0.73 | 608 | -0.021 | 0.468 | 1 |
| Bilateral | FMAJ | -1.25 | 631 | -0.043 | 0.212 | 1 |
|  | FMIN | 0.55 | 616 | 0.015 | 0.582 | 1 |
| Right | ATR | 0.08 | 629 | 0.003 | 0.935 | 1 |
|  | CAB | -1.53 | 643 | -0.055 | 0.127 | 1 |
|  | CCG | -1.36 | 629 | -0.049 | 0.174 | 1 |
|  | CST | -1.45 | 630 | -0.055 | 0.147 | 1 |
|  | ILF | -2.01 | 629 | -0.073 | 0.045 | 0.802 |
|  | SLFP | 0.76 | 625 | 0.028 | 0.446 | 1 |
|  | SLFT | -0.16 | 606 | -0.006 | 0.875 | 1 |
|  | UNC | -0.71 | 615 | -0.026 | 0.480 | 1 |

Associated *t*-values, degrees of freedom, standardized beta coefficients, uncorrected *P*-values, and interaction with tract *P*-values for impairment.

**Table S4.**
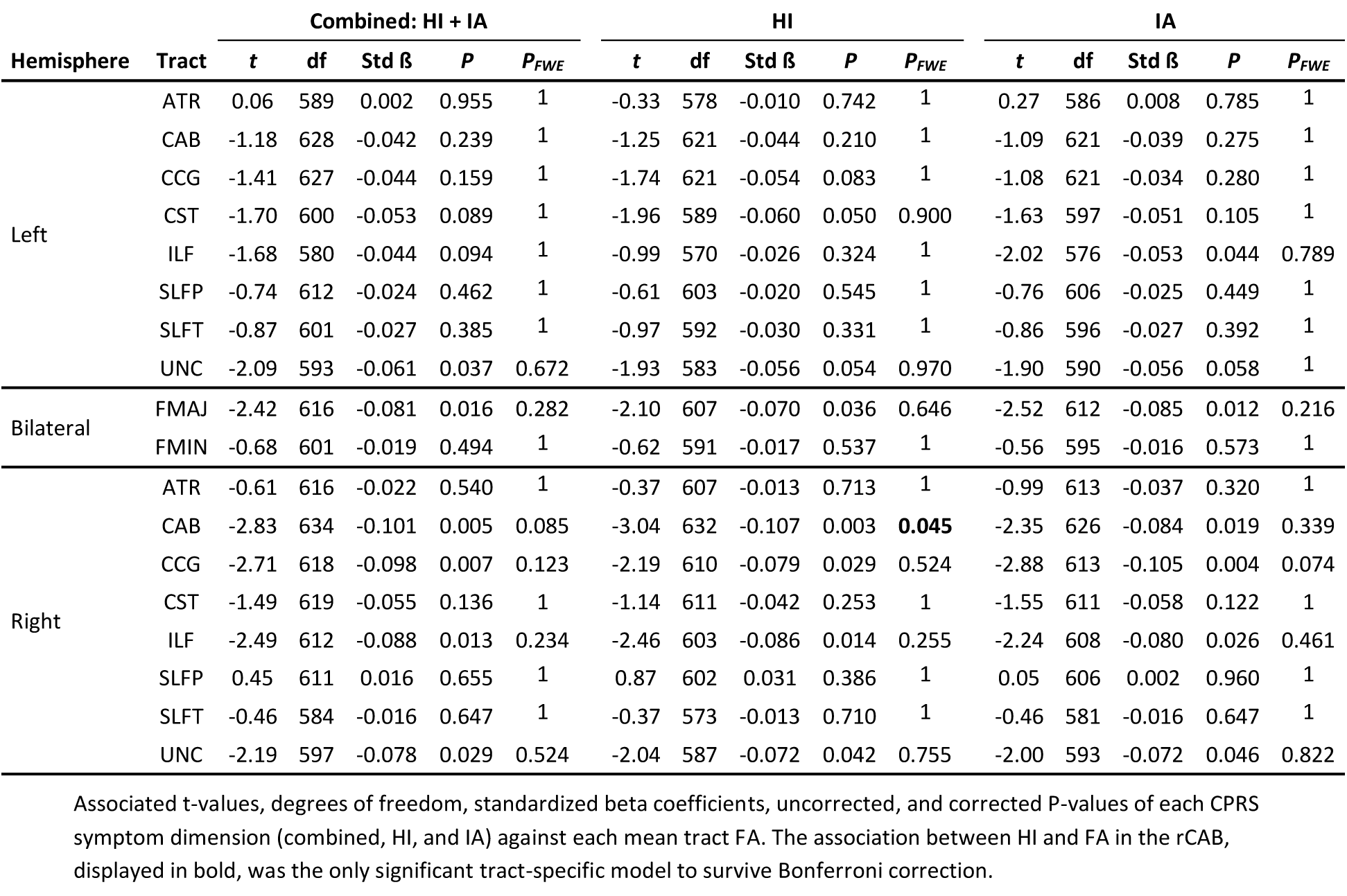
Symptom dimension score tract-specific model analyses.

| Hemisphere | Tract | Combined: HI + IA |  |  |  |  | HI |  |  |  |  | IA |  |  |  |  |
| --- | --- | --- | --- | --- | --- | --- | --- | --- | --- | --- | --- | --- | --- | --- | --- | --- |
| | | *t* | *df* | Std $\beta$ | *P* | *P<sub>FWE</sub>* | *t* | *df* | Std $\beta$ | *P* | *P<sub>FWE</sub>* | *t* | *df* | Std $\beta$ | *P* | *P<sub>FWE</sub>* |
| Left | ATR | 0.06 | 589 | 0.002 | 0.955 | 1 | -0.33 | 578 | -0.010 | 0.742 | 1 | 0.27 | 586 | 0.008 | 0.785 | 1 |
|  | CAB | -1.18 | 628 | -0.042 | 0.239 | 1 | -1.25 | 621 | -0.044 | 0.210 | 1 | -1.09 | 621 | -0.039 | 0.275 | 1 |
|  | CCG | -1.41 | 627 | -0.044 | 0.159 | 1 | -1.74 | 621 | -0.054 | 0.083 | 1 | -1.08 | 621 | -0.034 | 0.280 | 1 |
|  | CST | -1.70 | 600 | -0.053 | 0.089 | 1 | -1.96 | 589 | -0.060 | 0.050 | 0.900 | -1.63 | 597 | -0.051 | 0.105 | 1 |
|  | ILF | -1.68 | 580 | -0.044 | 0.094 | 1 | -0.99 | 570 | -0.026 | 0.324 | 1 | -2.02 | 576 | -0.053 | 0.044 | 0.789 |
|  | SLFP | -0.74 | 612 | -0.024 | 0.462 | 1 | -0.61 | 603 | -0.020 | 0.545 | 1 | -0.76 | 606 | -0.025 | 0.449 | 1 |
|  | SLFT | -0.87 | 601 | -0.027 | 0.385 | 1 | -0.97 | 592 | -0.030 | 0.331 | 1 | -0.86 | 596 | -0.027 | 0.392 | 1 |
|  | UNC | -2.09 | 593 | -0.061 | 0.037 | 0.672 | -1.93 | 583 | -0.056 | 0.054 | 0.970 | -1.90 | 590 | -0.056 | 0.058 | 1 |
| Bilateral | FMAJ | -2.42 | 616 | -0.081 | 0.016 | 0.282 | -2.10 | 607 | -0.070 | 0.036 | 0.646 | -2.52 | 612 | -0.085 | 0.012 | 0.216 |
|  | FMIN | -0.68 | 601 | -0.019 | 0.494 | 1 | -0.62 | 591 | -0.017 | 0.537 | 1 | -0.56 | 595 | -0.016 | 0.573 | 1 |
| Right | ATR | -0.61 | 616 | -0.022 | 0.540 | 1 | -0.37 | 607 | -0.013 | 0.713 | 1 | -0.99 | 613 | -0.037 | 0.320 | 1 |
|  | CAB | -2.83 | 634 | -0.101 | 0.005 | 0.085 | -3.04 | 632 | -0.107 | 0.003 | **0.045** | -2.35 | 626 | -0.084 | 0.019 | 0.339 |
|  | CCG | -2.71 | 618 | -0.098 | 0.007 | 0.123 | -2.19 | 610 | -0.079 | 0.029 | 0.524 | -2.88 | 613 | -0.105 | 0.004 | 0.074 |
|  | CST | -1.49 | 619 | -0.055 | 0.136 | 1 | -1.14 | 611 | -0.042 | 0.253 | 1 | -1.55 | 611 | -0.058 | 0.122 | 1 |
|  | ILF | -2.49 | 612 | -0.088 | 0.013 | 0.234 | -2.46 | 603 | -0.086 | 0.014 | 0.255 | -2.24 | 608 | -0.080 | 0.026 | 0.461 |
|  | SLFP | 0.45 | 611 | 0.016 | 0.655 | 1 | 0.87 | 602 | 0.031 | 0.386 | 1 | 0.05 | 606 | 0.002 | 0.960 | 1 |
|  | SLFT | -0.46 | 584 | -0.016 | 0.647 | 1 | -0.37 | 573 | -0.013 | 0.710 | 1 | -0.46 | 581 | -0.016 | 0.647 | 1 |
|  | UNC | -2.19 | 597 | -0.078 | 0.029 | 0.524 | -2.04 | 587 | -0.072 | 0.042 | 0.755 | -2.00 | 593 | -0.072 | 0.046 | 0.822 |

Associated *t*-values, degrees of freedom, standardized beta coefficients, uncorrected, and corrected *P*-values of each CPRS symptom dimension (combined, HI, and IA) against each mean tract FA. The association between HI and FA in the rCAB, displayed in bold, was the only significant tract-specific model to survive Bonferroni correction.

**Table S5.**
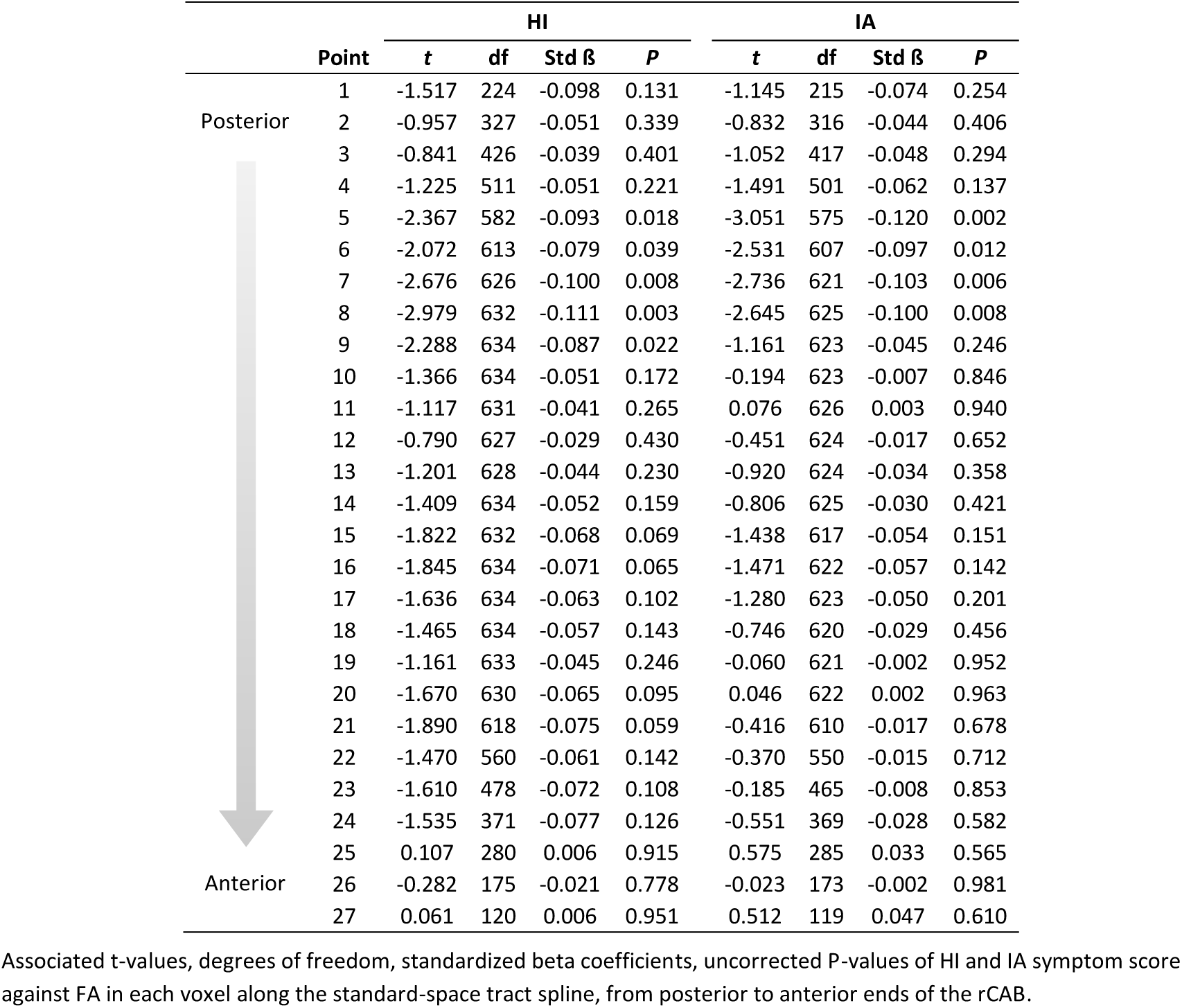
Within-tract model analysis: Point-wise comparison of FA along rCAB.

|  | Point | HI |  |  |  | IA |  |  |  |
| --- | --- | --- | --- | --- | --- | --- | --- | --- | --- |
| | | t | df | Std $\beta$ | P | t | df | Std $\beta$ | P |
| Posterior | 1 | -1.517 | 224 | -0.098 | 0.131 | -1.145 | 215 | -0.074 | 0.254 |
|  | 2 | -0.957 | 327 | -0.051 | 0.339 | -0.832 | 316 | -0.044 | 0.406 |
|  | 3 | -0.841 | 426 | -0.039 | 0.401 | -1.052 | 417 | -0.048 | 0.294 |
|  | 4 | -1.225 | 511 | -0.051 | 0.221 | -1.491 | 501 | -0.062 | 0.137 |
|  | 5 | -2.367 | 582 | -0.093 | 0.018 | -3.051 | 575 | -0.120 | 0.002 |
|  | 6 | -2.072 | 613 | -0.079 | 0.039 | -2.531 | 607 | -0.097 | 0.012 |
|  | 7 | -2.676 | 626 | -0.100 | 0.008 | -2.736 | 621 | -0.103 | 0.006 |
|  | 8 | -2.979 | 632 | -0.111 | 0.003 | -2.645 | 625 | -0.100 | 0.008 |
|  | 9 | -2.288 | 634 | -0.087 | 0.022 | -1.161 | 623 | -0.045 | 0.246 |
|  | 10 | -1.366 | 634 | -0.051 | 0.172 | -0.194 | 623 | -0.007 | 0.846 |
|  | 11 | -1.117 | 631 | -0.041 | 0.265 | 0.076 | 626 | 0.003 | 0.940 |
|  | 12 | -0.790 | 627 | -0.029 | 0.430 | -0.451 | 624 | -0.017 | 0.652 |
|  | 13 | -1.201 | 628 | -0.044 | 0.230 | -0.920 | 624 | -0.034 | 0.358 |
|  | 14 | -1.409 | 634 | -0.052 | 0.159 | -0.806 | 625 | -0.030 | 0.421 |
|  | 15 | -1.822 | 632 | -0.068 | 0.069 | -1.438 | 617 | -0.054 | 0.151 |
|  | 16 | -1.845 | 634 | -0.071 | 0.065 | -1.471 | 622 | -0.057 | 0.142 |
|  | 17 | -1.636 | 634 | -0.063 | 0.102 | -1.280 | 623 | -0.050 | 0.201 |
|  | 18 | -1.465 | 634 | -0.057 | 0.143 | -0.746 | 620 | -0.029 | 0.456 |
|  | 19 | -1.161 | 633 | -0.045 | 0.246 | -0.060 | 621 | -0.002 | 0.952 |
|  | 20 | -1.670 | 630 | -0.065 | 0.095 | 0.046 | 622 | 0.002 | 0.963 |
|  | 21 | -1.890 | 618 | -0.075 | 0.059 | -0.416 | 610 | -0.017 | 0.678 |
|  | 22 | -1.470 | 560 | -0.061 | 0.142 | -0.370 | 550 | -0.015 | 0.712 |
|  | 23 | -1.610 | 478 | -0.072 | 0.108 | -0.185 | 465 | -0.008 | 0.853 |
|  | 24 | -1.535 | 371 | -0.077 | 0.126 | -0.551 | 369 | -0.028 | 0.582 |
|  | 25 | 0.107 | 280 | 0.006 | 0.915 | 0.575 | 285 | 0.033 | 0.565 |
| Anterior | 26 | -0.282 | 175 | -0.021 | 0.778 | -0.023 | 173 | -0.002 | 0.981 |
|  | 27 | 0.061 | 120 | 0.006 | 0.951 | 0.512 | 119 | 0.047 | 0.610 |

Associated t-values, degrees of freedom, standardized beta coefficients, uncorrected P-values of HI and IA symptom score against FA in each voxel along the standard-space tract spline, from posterior to anterior ends of the rCAB.

**Table S6.**
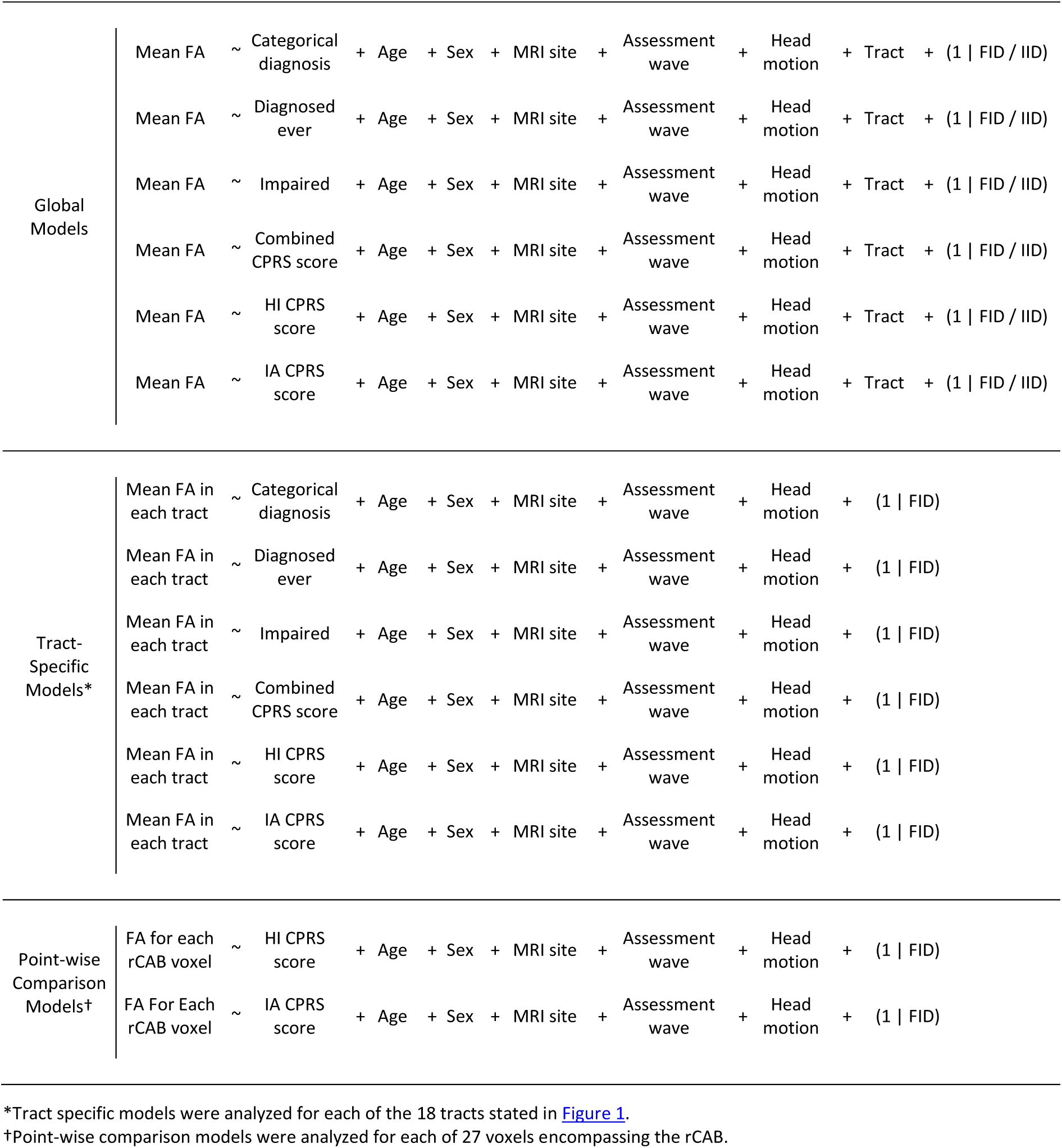
All linear mixed effects regression models used in all analyses.

